# Separated attractors in neural landscape of motor cortex encoding motor learning

**DOI:** 10.1101/2024.10.01.611949

**Authors:** Xuanting Liu, Yanzi Wu, Xiahan Ru, Rongrong Li, Ke Si, Wei Gong

## Abstract

Animals gain motor learning via decrease of variation through repeated training. The secondary motor (M2) cortex shows an indispensable role in the learning process of the rotarod-learning task. Yet, it remains unclear how population decoding in M2 cortex guides the repetitive training to transform into motor enhancement. We recorded neuronal population activity using Ca2+ imaging during this enhancement revealing that neuronal population correlates of the persistent internal learning state evolves in the process of motor learning. With the behavioral micro-states analysis, we identify the growing periodicity, stability, and consistency with two gradually clearer point attractor in the M2 neural state space. The results show the evolution of attractors in M2 participate in decrease of training-acquisition behavior variation and provide a general framework for the mapping between arbitrary non-task motor learning and neural topological structure.

## Introduction

The accelerating rotarod-learning task is a widely used rodent motor learning paradigm, in which animals learn steady stepping movements on a rotating rod over training[1–3]. In the learning paradigm, motor learning often means reducing motor variability[4]. At the beginning of the learning paradigm, motor variability can be construed as a means of exploring motor space. When expert phase, less motor variability improves performance and reduces costs, thus steering the motor system towards steady and robust patterns[5].

Currently, “time latency” is used as the assessment criterion for the rotarod learning task in mice, though this approach has shown limited effectiveness. While some studies employ multi-parameter assessments or focus on individual body parts to further evaluate rotarod performance, a detailed behavioral analysis of traditional rotarod motion is still lacking[6, 7]. Without refined analysis, it remains challenging to deeply understand the learning process, learning strategies, key actions, and corresponding neural control modules.

Motor learning involves coordinated activities in motor cortex[7, 8]. The lesions or inactivation of motor cortex has been shown to impair motor learning[7]. Motor cortex has been suggested to be necessary for progressing steady and robust motor movements in mice. In the motor system, it has been shown that motor cortex demonstrates global changes in motor learning[9, 10]. However, an important open question is how the activity of these neural subpopulations during behavior executes collaborative functions. The traditional averaging method often obscures individual cell activity patterns, and consequently, relatively few single-unit recordings have been conducted. As a result, a critical gap is the lack of detailed information regarding the population-collaboration activity of neural subpopulations during the process of motor learning.

Ca2+ imaging with single-cell resolution is enabled in moving animals[11, 12]. Application of this approach to the secondary motor(M2) cortex has shown that the statistical measurement of cells movement response in M2 is essential for motor learning[13]. However, there is no clear direct linear coding evidence between single-unit spiking patterns and behavior. In other system, modeling neural populations as dynamical systems has revealed latent correlation between dynamics of population activity and behavior. Here, we employ miniature head-mounted microscopes and describe the attractor in M2 cortex. Our findings reveal that the M2 cortex exhibits an additional attractor in rotarod task compared to natural state. This second attractor appears crucial in reducing motor variability.

## Result

### Internal representation of space in the rotarod training

To track the formation and evolution of neural ensembles in the M2 cortex of awake, head-fixed mice across motor learning, we used a custom miniScope system to record the calcium activity of M2 cortex while mice learned a rotarod task (Fig. 1A). The rotarod task consisted of a learning phase of 5 days. For relatively stable recording, each training day was divided into two states: testing state, continuous rotarod task with miniScope, and training state, accelerating rotarod task to train motor ability (Fig. 1B). We trained several mice on the task, all reaching effective improvement of performance score (Supplementary Fig. 1A) within 5 days. Based on this, the last day was defined as expert phase.

**Figure 1.**
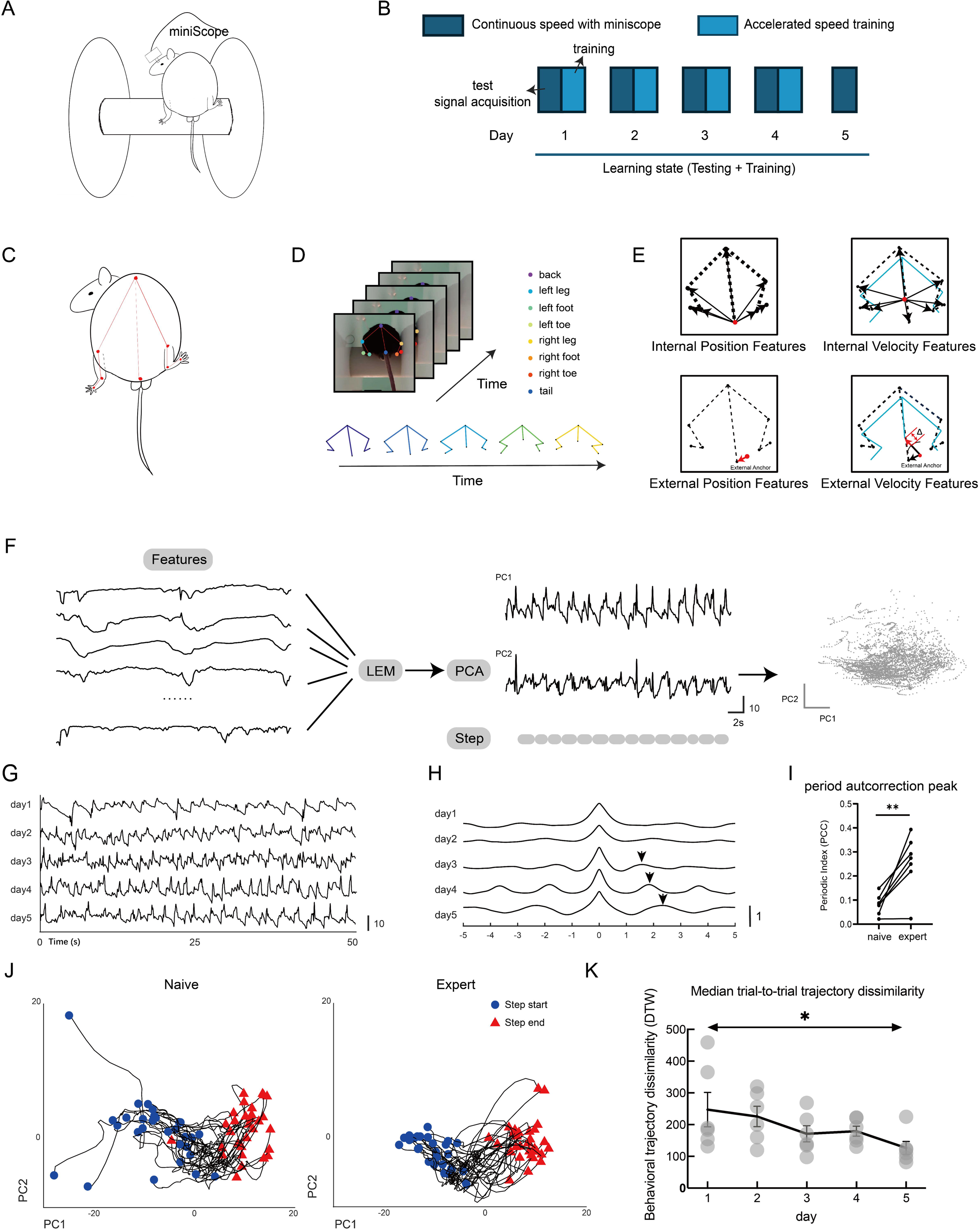
Rotarod learning and behavioral pose estimation. (A) Rotarod experimental setup, mouse was equipped with a head-mounted micro microscope placed on a slowly rotating rod, baffles were set on both sides of the mouse to limit the space. (B) Training regimen, each day consisted of continuous speed test and accelerated speed training. (C) Pose estimation, eight anatomical landmarks were selected for tracking: the back, tail, left and right legs, left and right feet, and left and right toes. (D) Video tracking, colored dots denote body part labels. Body part positions across successive frames form a posture sequence. (E) Behavioral feature extraction. Internal features extraction: point-to-point distances between body parts within each frame (N = 28 with 8 body parts) and speed of each point of the body relative to the tail in the adjacent frame (N = 64 with 8 body parts). External feature extraction: distances between body parts and external anchors within each frame (N = 8 with 8 body parts) and speed of each point of the body relative to the tail in the adjacent frame (N = 8 with 8 body parts), each external anchor is the center of motion range of each body part throughout the video. (F) Diagram of dimensionality reduction. Primary nonlinear features were mapped to 6 dimensions by Laplacian Eigenmaps (LEM), and then feature vectors were mapped to a two-dimensional plane by Principal Component Analysis (PCA). First two PC accounts for 68.8% ± 2.0% of the total variance of behavior activity (N = 6 mice). (G) PC1 of the behavioral space of mice over a 50-second window for each day, showing a pattern of reciprocating movement. (H) Autocorrelation function graph of PC1 during training period. A peak gradually appeared around 2 seconds with training days increased. (I) Periodic index of period autocorrelation peak over learning, measured by the height and position of the first non-central peak relative to central peak. (p < 0.01, n = 7 mice, regression, mean ± S.E.M.) (J) Example snippets of behavior trajectories of mice’s steps in low-dimensional space across learning for one mouse. Each line corresponds to a single trial, beginning at step start (circle markers) and ending at step end (triangle markers). (K) Stabilization of the mouse behavior across learning. X axis shows learning days. The median pairwise distance between trajectories on different trials (measured by dynamic time warping algorithm) is plotted as a black line (± SEM) with individual mouse means shown as gray circles.

Quantifying motor skill learning by the time of rotarod latency is useful in studying the motor learning performance, and the technique remains widely used in the field[14, 15]. However, this approach does not allow the identification of all evolutionary process in the motor learning [16–20] that may be relevant to interpreting the activity of M2 neurons, making it difficult to quantify the transient response of motor learning. To overcome this limitation, we constructed internal and external behavioral features into a behavioral space. In order to accurately observe the movement of mice on the rotarod, we selected the back of mice as the observation direction, and used the back, knee joint, ankle, toe and tail root of mice as marker point (Fig. 1C). We first used DeepLabCut [21] to monitor the position and posture of a mouse throughout video frames (Fig. 1D). Then we extract the mutual distance and speed between body parts location as internal features, and the distance and speed from body position at any given time to the average as external features (Fig. 1E).

To construct the relationships among the feature ensemble activity patterns, we sought to embed the data in a reduced-dimensional space of behavioral activity. Non-linear dimensionality reduction approaches enable such embedding, making them more suitable than linear methods in many cases [20, 22]. Here, we apply the non-linear dimensionality reduction algorithm Laplacian Eigenmaps[23] (LEM) to the behavior vectors, embedding abstract behavior process into a low-dimensional space. With this strategy, the high-dimensional, continuous, and time-varying behavioral series can be decomposed to various quantifiable movement parameters and low-dimensional behavior map. Then, to visualize the properties more easily, we plotted the space in 2D using principal component analysis (PCA) (Fig. 1F). PCA indicated that the first two PC accounts for 68.8% ± 2.0% of the total variance of behavior activity (Supplementary Fig. 1B) (N = 7 mice). Standard behavior is mostly confined to a concentrated range in the space, while corrective behavior performs a shifting away from the range. Specifically, PC1 captured major information of behavior activity, which indicates the major motion period. The trajectories in a concentrated range in PC1 exhibits a reciprocating pattern indicating the roughly same motion period (Fig. 1G).

It has been revealed that motor learning decreases the variability of behavior[4]. Therefore, we ask the particular paraments, periodicity and consistency, of variability within the motor learning process. To investigate the possible periodicity, we employed the autocorrelation methods, often used in curve characteristic analysis, to construct the periodograms in PC1[24] (Fig. 1H). We found that within training, a peak around 2 seconds emerged in autocorrelation curve (Fig. 1H), and the height of peak increased as training going on (Fig. 1I), which means the behavior in the major dimension shows similarity with the behavior 2 seconds before or after. The similarity gets strengthened significantly when expert (Fig. 1I), indicating that the mice acquire rotarod sequences periodicity within the training.

The consistency is indicated by the trace in behavior space (Fig. 1J). In low-dimensional space, Mice’s similar steps in different scales caused by shooting angle noise were adjusted into adjacent trajectories, which means changes of trajectories in the low-dimensional space robustly represent the process of motor learning in real world (Supplementary Fig. 1C). Mice’s behavioral trajectories in low-dimensional space were more disorganized when naive and gradually became more organized during acquisition of rotarod training (Supplementary Fig. 1D). Furthermore, a mixed-effects ANOVA on the average trial-to-trial variability of mice’s behavioral trajectories across training days (Fig. 1J) revealed a significant effect of time (p < 0.05), indicating that the mice acquire rotarod sequences stabilization by the repetitive behavior period (Fig. 1K).

### Rotarod training adds an additional attractor in M2 neural space and displays long memory

The M2 cortex has been reported to be responsible for motor learning[7, 25, 26]. However, the population dynamics patterns of the M2 cortex during rotarod training have not been thoroughly investigated. To further characterize the activity patterns of neurons during motor learning, we used miniScope to track M2 spike neuron in awake, free-moving mice. Across five days training, approximate number but different individuals of neurons in M2 participate in the rotarod task (Supplementary Fig. 2A). Approximately half of the neuronal activity locations remained consistent between adjacent days (Supplementary Fig. 2B). For any individual cell, which participate in coding the behavior, no significant consistent coding patterns exists across different learning days (Fig. 2A).

**Figure 2.**
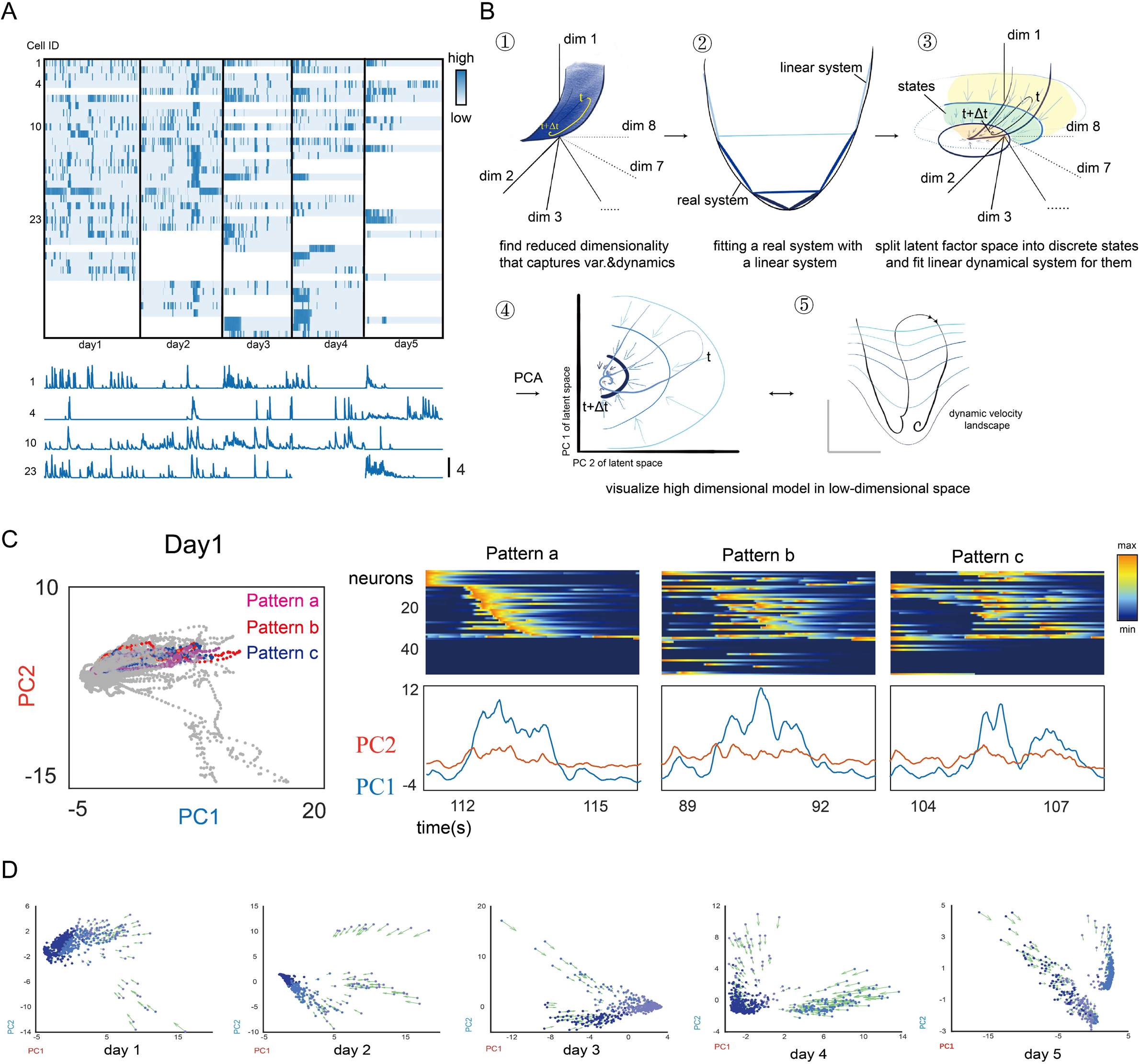
M2 contains a similar attractor structure. (A) Calcium trace of all cell over the training period. Clustering of recorded neurons in M2 cortex. (B) Schematic illustrating rSLDS analysis. Neural activity reveals an integrator dimension that correlates with behavior on rotarod task. (C) Release sequence of adjacent neural pattern in PCA space. Different neural pattern shared similar release sequence and fluctuation of PC1 and PC2. (D) Neural state space with population trajectories across learning episodes, colored by discrete state *z_t_*. The arrow shows the force *f_t_* for each continuous latent factor (*x*).

To uncover the neural population encoding patterns, we then employ a population analysis using a fitted recurrent switching linear dynamical system (rSLDS)[27]. The rSLDS model effectively simplifies the complex non-linear dynamics of neural systems, facilitating a more comprehensive understanding of neural encoding. This method initially transforms the recorded neural data into a set of latent variables (continuous variables). The latent variables, representative of neural state at any single moment, was on a continuous “terrain” and in a neural “gravitational field”, which means latent state would be captured by continuous attractors[28] (Fig. 2B ➀). To further simplify the system, a linear discrete model, in which latent variables obtain a discrete state depends on its location determining its terrain structure, was used to fit the real continuous systems (Fig. 2B ➁). Through this approach, rSLDS split latent factor space into discrete states and constitute the entire dynamic space (Fig. 2B ➂). Each discrete state is described by a unique dynamic matrix that decide where it charge and a transition matrix that decides how neural activity evolves over time. In this way, rSLDS reveal the dynamical properties of the neural circuit.

Finally, we utilize principal component analysis (PCA) (Fig. 2B ➃) to visualize the high-dimensional model in a low-dimensional space, providing an intuitive representation of the system’s dynamics. In single day, adjacent neuron patterns show similar release sequences along the trajectories (Fig. 2C). While from the whole training time-scale, vectors of 2D flow field in PC indicate how neural dynamics evolve according to the fit rSLDS model (Fig. 2D). This revealed one or two regions of low vector flow that forms an approximate point attractor, meaning that the neural population activity vector tends to move towards one or two points. Meanwhile, the main attractor performs a similar dynamic structure (Supplementary Fig. 2E), as well as its consistent stability (Supplementary Fig. 2F).

We are interested in the collective principal activity characteristics of neurons. To figure out the characteristic of neural activity, autocorrelation analysis is employed. Another use of autocorrelation coefficient is considered as a measure of temporal stability of the neuronal activity[29]. “Decoupling time”, as the time it takes to descend from the central crest to the zero, which means the shortest time for the curve to become orthogonal to itself after time *t*, represents the temporal stability. We calculate the autocorrelation coefficient and decoupling time of PC1 from different days (Fig. 3A). The decoupling time increases with the learning process (Fig. 3B), while the decoupling time of expert neural activity significantly increased (Fig. 3C), indicating that the neural activity of mice in rotarod task was increasingly strengthened. What’s more, the performance of rotarod task is significantly explained this character of neural principal activity (Fig. 3D), which indicates the overall characteristics of M2 cortex govern the motor learning performance of mice.

**Figure 3.**
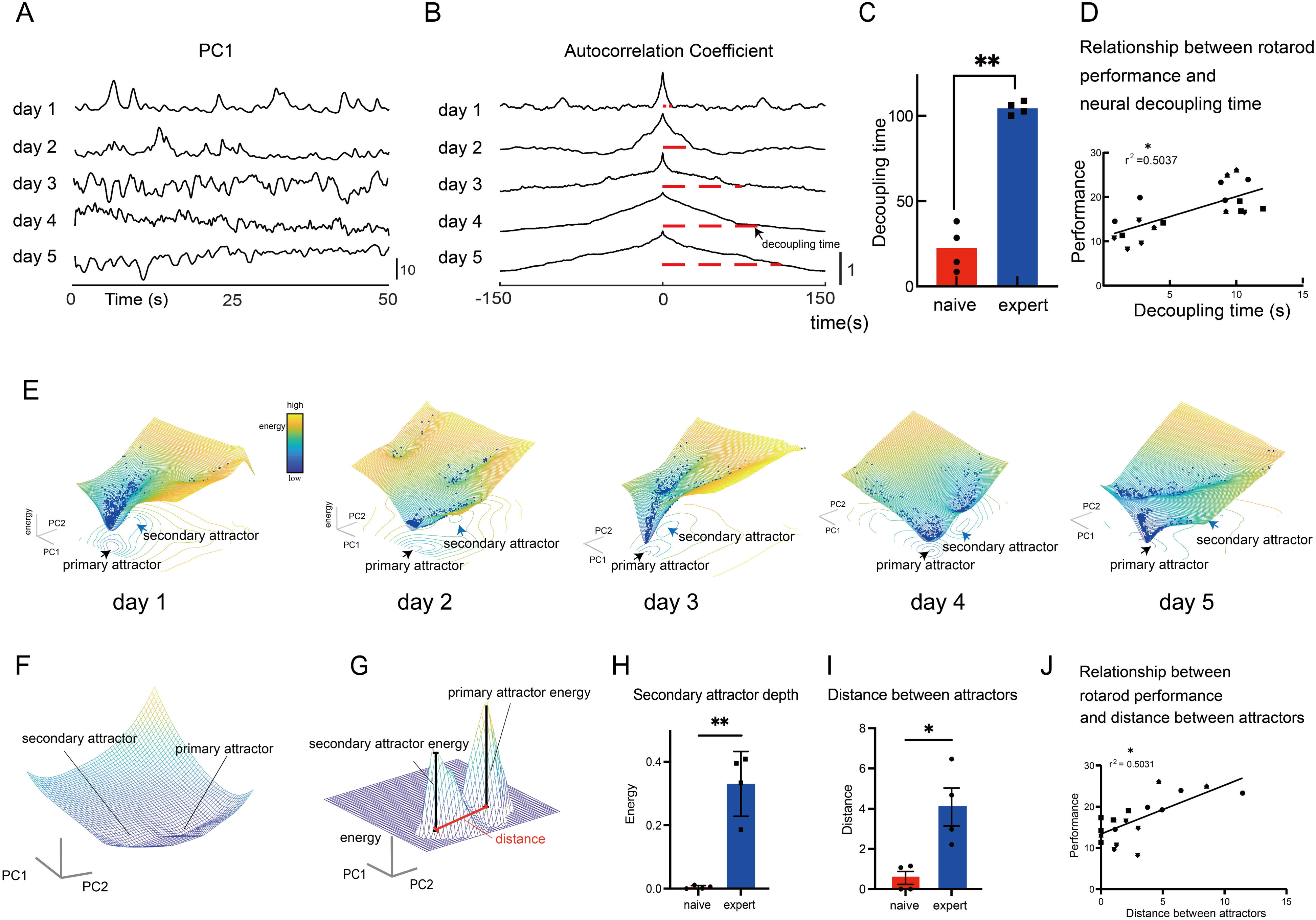
The strengthened temporal stability and more separated attractors along with motor performance improvement. (A) Example of snippet of PC1 in neural space over a 50-second window across learning for one mouse. (B) Autocorrelation coefficient of PC1, which is shown in A, in neural space across learning. Decoupling time is defined as the shortest time for the curve to become orthogonal to itself. (C) Decoupling time for autocorrelation analysis shown in B of expert or naïve phase (n = 4 mice, p<0.01, paired t-test). (D) Decoupling time shown in C versus corresponding rotarod performance of individual mice (r^2^ = 0.507, n = 4 mice; asterisk indicates P < 0.05, regression). (E) Representative neural dynamics landscape of a single mouse across five training days, with attractors integrated by 2D flow field. (F) Representative 3D landscape for two attractors. (G) Inferred depth of primary and secondary attractors from Flood-Fill algorithm and distance from dynamic fixed-point in neural dynamics landscape shown in F. (H) Depth of secondary attractor across training in naïve or expert state (p<0.01, n = 4 mice, paired t-test). (I) Distance between two attractors across training (p<0.05, n = 4 mice, paired t-test). (J) Relationship between performance and distance between attractors for individual mouse (r^2^ = 0.501, n = 4 mice; asterisk indicates P < 0.05, regression).

To visualize the manifold of the rSLDS model, we quantified the potential energy to construct 3D landscape. We integrated the flow-field vectors over distance in neural state space to get the energy at each point. Then we convert the energy into the height (z axis) of the landscape (Fig. 3E); the x-y axes are still represented by PC1 and PC2. In this topographic representation, a point attractor would appear as a bowl of slow rate of change at the base of a cone, reflecting the stable pattern emitting over a period.

The evolution of manifold is another noteworthy change within motion learning. To our surprise, when in the rotarod task, even in the naïve state, an additional point attractor tends to exist, meaning that the neural population activity vector tends to move toward another stable activity pattern (Fig. 3E). To measure the relative energy between two attractors (Fig. 3F), we introduce Flood Fill Algorithm to fill the “valley” in landscape. We then utilize the residual to calculate relative energy of two attractors and distance between attractors as well (Fig. 3G). Comparing the expert state to the naïve state, we discover that the additional point attractor was significantly deepened within motor learning (Fig. 3H), as well as the distance between attractors (Fig. 3I), which indicates an acquired and separated pattern emerged during training progresses. Furthermore, it shows that rotarod performance and distance between attractors are positively correlation during training phase, which means motor ability is significantly explained by gradually separated attractors (Fig. 3J).

### Separated neural pattern charging behavioral variability in M2 cortex

In the expert stage, it has been established that the additional point attractor is deepened (Fig. 3H), with the structure of the two attractors becoming increasingly distinct (Fig. 3I). The neuronal space will gradually form two separate regions corresponding to these attractors (Fig. 4A). When neuronal activity is captured by different attractors, the mice show different discreteness in their behavior, manifesting as two areas with different ranges in the behavioral space (Fig. 4B). Furthermore, the firing patterns of neurons differ between the two attractors (Fig. 4C), indicating a collective separation pattern that is commonly observed among expert of motor learning.

**Figure 4.**
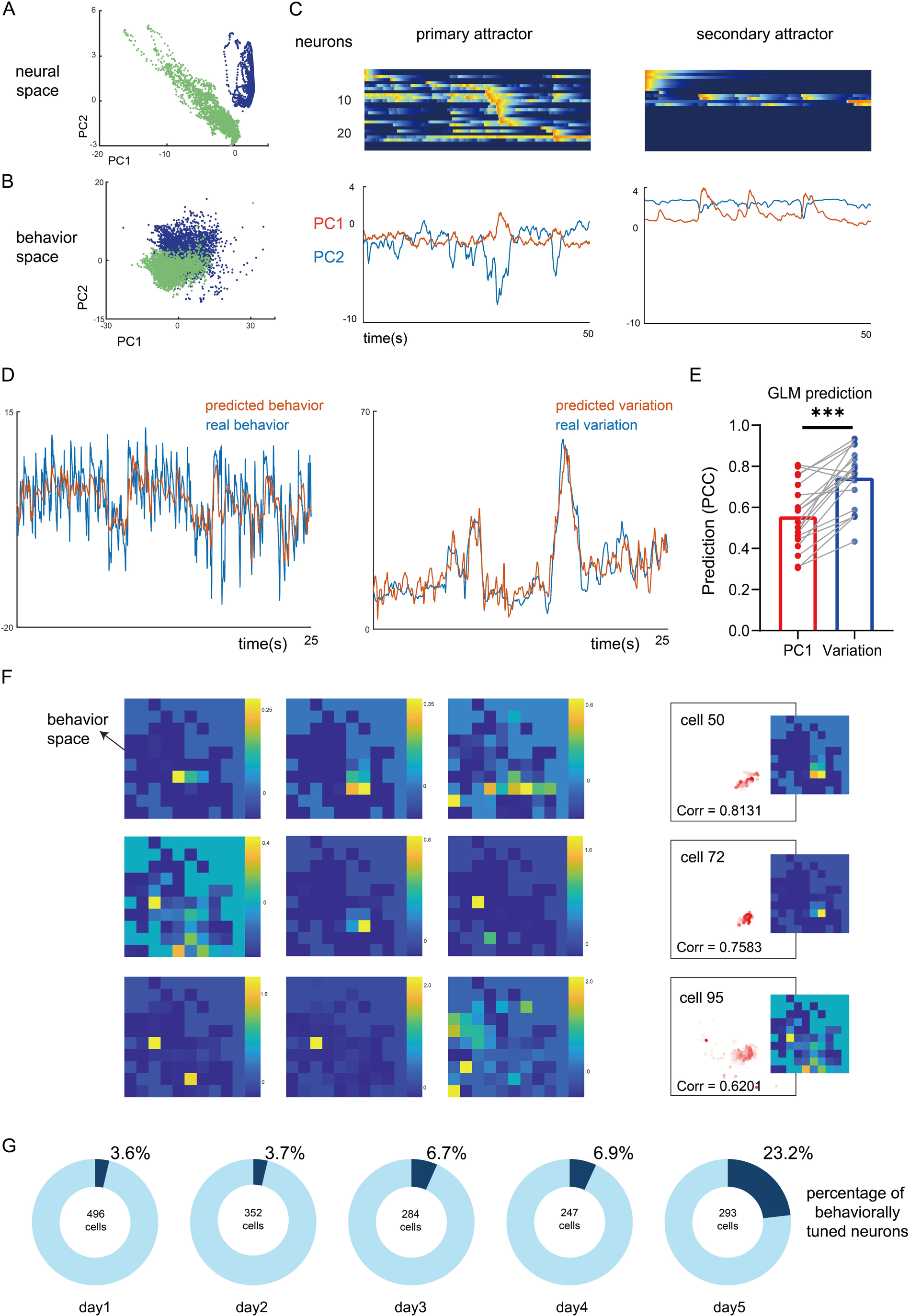
Mapping of Neuronal Regional Characteristics to Behavioral Regional Characteristics. (A) Neural state space with two separated population distribution. (B) When neural activity is attracted by two different attractors, the distribution range of behavioral features in the behavioral low-dimensional space is different. The colors match Figure A (C) Sequential activity of M2 neurons when captured by the primary attractor (left) and secondary attractors (right). (D) Real behavioral PC1 together with neural projections onto the weights that best predict behavioral PC1 (left), real behavioral variation together with neural projections onto the weights that best predict the behavioral variation(right). (E) Prediction from activity of neuronal populations in M2 cortex, based on Pearson’s correlation coefficient (PCC) between real and predicted behavioral PC1 and variation. The significant prediction differences for behavior PC1 and variation are shown (p<0.001, n = 4 mice, paired t-test). (F) Heat map represents the firing location of neurons in the behavioral low-dimensional space. A high color-bar value indicates a high average firing intensity of neurons in the unit behavior space. (G) Percentage of cells that have a high rate of firing in a particular behavioral area (five pies represent all the cells in each day).

To investigate the encoding of behavioral variability at the population level, a generalized linear model was employed to predict both behavioral PC1 and behavioral variability (Fig. 4D) using calcium activity data from the M2 cortex while mice were learning a rotarod task. The Pearson correlation coefficient between the predicted and actual behavioral PC1 and variability was used as a measure of predictive power. When the predictive power in both conditions was measured, the neuronal calcium signals demonstrated significantly greater predictive power for behavioral variability compared to behavioral PC1 (Fig. 4E). This suggests that, compared to the behavior itself, the information contained in neural energy more effectively accounts for behavioral variability, as a further explanation of neural pattern charging behavioral variability.

To further determine the limiting effect of cells on behavioral variability, a method similar to “Place Cell” is used to search for behavior-place-specific cells. Behavioral features were mapped onto a 2-dimensional PCA space, dividing it into uniformly sized micro-behavioral regions (Fig. 4F). By correlating neuronal firing times with the position of the mouse in this behavioral space, we were able to determine the specific behavioral state the neural was in at any given moment. To quantify the behavioral tuning of M2 neurons, an index of the correlation between the firing rate of each neuron and the activities within these micro-behavioral regions was calculated (Fig. 4F). Notably, certain neurons exhibited heightened firing intensity within a limited number of specific regions, as illustrated in Fig. 4F. These behaviorally tuned neurons were identified across all mice over a 5-day period. The proportion of these specialized cells, relative to the total population of neurons observed per day, increased progressively with motor learning (Fig. 4G). This suggests that the acquisition of behavior recruits more behavior-place-specific cells, which indicates more cells confined to a specific region reduce their variability in the behavioral space.

Furthermore, the direct correlation between neural energy and behavioral variability was detected. In the naive state, the secondary attractor is not prominent (Fig. 5A), with neuronal activity primarily captured by the primary attractor, although occasional upward jumps indicate an escape tendency from the primary attractor. However, in the expert state, two relatively independent attractors emerge (Fig. 5B). It was previously found that, after dimensionality reduction, the primary component (PC1) was found to capture the essential features of behavioral activity (Fig. 5B). Autocorrelation analysis revealed a behavioral periodicity of approximately 2 seconds (Fig. 1H), aligning with the time required for a single stride in mice, which means relatively consistent peak-valley period (Fig. 5C) runs through the learning process, whether it exhibits similarity as a “periodicity” or not. This behavior variability from periodicity can be explained by variance, where high periodicity means low variance. The sliding variance of behavior in a 2.5 seconds window (Fig. 5D) is used to evaluate the variability in behavior. Obviously, the neural energy did not exhibit similar periodicity same as behavior (Fig. 5E). When neuronal activity was captured by the primary attractor, neural energy displayed peaks approximately every 10 seconds (Fig. 5E), while captured by the secondary attractor, the neural energy remained relatively stable at a higher level. This misalignment of cycles denies M2’s direct encoding of behavior itself, but the consistency of the fluctuation characteristics of behavioral variance to neuron suggests potential neural energy activity matching (Fig. 5D). Surprisingly, when behavioral variability reaches its peak, an increase in neuronal energy has been shown to play a role, actively reducing the fluctuations in behavior to a more controlled and lower level (Fig. 5F). This phenomenon often occurs at the first attractor no matter the mice are in naive state or expert state. We infer there is a regulatory mechanism wherein heightened neuronal activity within the M2 cortex, contributes to maintaining behavioral consistency by counteracting excessive variability.

**Figure 5.**
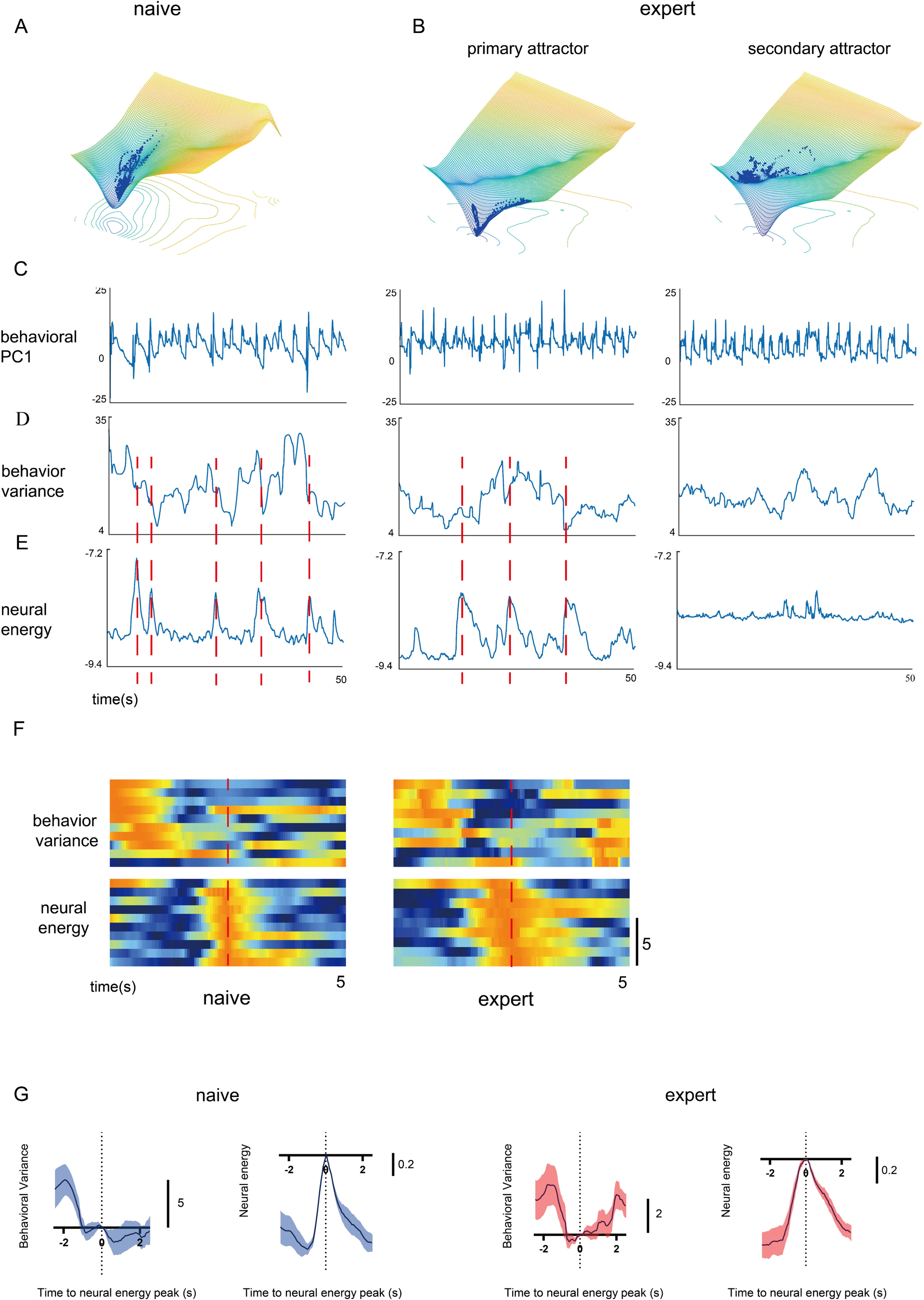
Neural energy fluctuation inhibits behavioral variability. (A) Inferred 3D dynamic neural landscape in naive mouse with population distribution. (B) Inferred 3D dynamic neural landscape in expert mouse with population distribution captured by the primary attractor (left) and secondary attractor (right). (C) Behavioral PC1, behavioral variability, and neuronal energy curves of the same duration (50s) when mice are naive (left), captured by the primary attractor in expert state (middle) and captured by the secondary attractor in expert state (right). (K) Heat maps of behavioral variability and neural energy with warm colors indicating higher values. (L) Statistical plots of all behavioral variability and nerve energy over the same period of time (5s) after aligning nerve energy peaks.

Next, the mean activation of each neural energy peak was computed. Regardless of whether in the naive or expert state, the peaks in neural energy coincided with troughs in behavioral variance (Fig. 5G). This finding suggests the presence of a specific neuronal firing pattern within the M2 cortex that exerts a modulatory effect, potentially reducing behavioral variability. It indicates an inhibitory effect of neural energy on behavioral variability.

## Discussion

In this study, a detailed behavioral analysis was employed to construct a behavioral space for mouse rotarod performance (Fig. 1A-F). Detailed behavioral analysis is essential, particularly in studies focusing on context-dependent behaviors, micro-behaviors, or behavioral progressions that are difficult to observe visually. In previous studies[30], behavior learning was often sequential, meaning that a stimulus (or not) goes ahead and then the behavior follows, usually with a zero-moment used to locate the starting point of the behavior. This means that at least three parts have been introduced when learning: understanding instructions, learning behavioral sequences, and muscle training. At the same time, the standard stimulation of mouse behavior suppressed its exploration, making it an “obedient machine” rather than a “boy learning to ride a bicycle”. Even when behavioral models bifurcate, they are usually limited and used to study behavioral decisions with low variation. However, motor learning is different from working memory, emphasizing the acquisition of a skill, which means learning path is more concerned[4]. Learning is often path-dependent, where a small number of correct attempts at the beginning will become the cornerstone of its subsequent exploration. This exploration path is unreachable in previous model construction because of its binarization of “acquired” or “unacquired”. A refined behavioral analysis can capture these subtle nuances, allowing for a more comprehensive understanding of motor processes[18, 31]. Thus, the spontaneous or autonomous behavior of animals can be better studied.

It was observed that during the rotarod learning process, the behavior of mice in the expert stage exhibited periodicity and consistency (Fig. 1G-K), showing a decrease in behavioral variability. This indicates that, with motor learning, the system transitions toward stable and robust motor patterns. Recent studies have further suggested that motor variability may be a critical feature of how sensorimotor systems function and learn[4]. Introducing the concept of motor variability provides a deeper understanding of motor learning. Variability in motor actions is inevitable, as no matter how much we practice, it is difficult to replicate identical actions perfectly. This is partly because our nervous system is inherently ‘noisy’, meaning that even well-practiced movements are subject to variation[32, 33]. Importantly, reductions in motor variability are often associated with improved task performance and skill acquisition, as noted in previous studies[4]. Thus, motor variability can serve as a valuable tool for evaluating mice performance during motor learning tasks, offering insights into their learning progress and the quantitative adaptability of their motor systems. By examining motor variability, we can better understand how well the motor learning process is unfolding, and how effectively the system is stabilizing toward more efficient movement patterns.

In this study, we utilized a recurrent Switching Linear Dynamical System (rSLDS) to describe the intrinsic neuronal space, revealing the gradual formation of a bistable attractor system (Fig. 2). Understanding how neuronal population activity in the M2 region encodes the rotarod learning task is fundamentally about mapping external spatial representations to intrinsic neuronal dynamics. Previous studies have demonstrated that neuronal intrinsic structures can map to external spaces, which may represent concrete physical environments, abstract behavioral, and emotional spaces. For example, it has been shown that the intrinsic activity space of individual grid cells forms a ring, where specific locations on the ring correspond to the animal’s position in a real environment[34]. In studies of motivation and decision-making during foraging, the Landscape Diffusion Model has been employed to construct an oscillatory dynamical space, reflecting the complexity of the animal’s behavior[35]. Additionally, researchers have built a line attractor-like neuronal space, where neuronal trajectories along this attractor were associated with increasing competitive behaviors, suggesting shifts in the animal’s emotional state[27].

Constructing a reasonable internal space can provide deeper insights into neuronal activity. For instance, neuronal activity in the auditory cortex has been effectively described by an internal structure derived through linear dimensionality reduction (PCA), which aids in understanding auditory input processing[36]. However, in many cases, structures obtained solely from linear dimensionality reduction approaches are insufficient to explain external behaviors[37]. Nonlinear dimensionality reduction techniques, such as Laplacian Eigenmaps (LEM)[37] or neural network[38, 39], have been applied to elucidate simpler motor behaviors. Yet, for more complex, spontaneous behaviors, these dimensionality reduction methods often lack interpretable neuronal structures. In contrast, the rSLDS[27] we employed incorporates dynamical concepts, focusing not only on the current position of neural activity within the internal structure but also on predicting its future trajectory (Fig. 2D). This approach allows for a more comprehensive understanding of neuronal population dynamics and their mapping to external spaces, thereby providing a more robust explanation of the processes underlying rotarod learning.

Energy is always a key issue of the construction of the energy landscape of neurons. In the attempt of the rSLDS dynamics constructor[27], they defined the length of vector as energy. We have a concern that it is an estimate of “kinetic energy” versus “potential energy”. Therefore, in our construction, we use the integral of the change in the discrete state determined by the inference in the real situation to evaluate the potential energy of the space (Fig. 3). Unavoidably, due to the possible rotational characteristics of linear dynamics, in extreme cases, the dynamics can perform circular motion without energy loss on a toroidal attractor, which makes rough estimate acceptable. However, it is inconsistent with the physical properties of neurons and rejects the possibility of using potential energy to infer latent motion. Due to this conflict, rSLDS will not be able to fully leverage its advantage of allowing discrete states estimation to segment regions.

On the other hand, rSLDS have another characteristic, which is the fixity of the energy landscape. rSLDS attributes the changes in the dynamic structure to discrete state switching over time, meaning that the changes in its dynamics over time can be captured and estimated. However, rSLDS attempts to attribute this change to the location of continuous latent variables, which will not be able to estimate the dynamic structure of variation over time. Changes in the energy landscape of thirst and hunger over time, as Luo[35] has done, will not be achievable using this type of estimation. We tried to put all the energy landscape data of training days into a single dynamic structure rSLDS estimation. However, considering the changes in the dynamics and unknown cell types, no interesting results were found. We believe that using linear dynamics to evaluate behavior will be a feasible way, considering the approximate invariance and proportional variability of behavior sequence. In subsequent research, we will use network methods to train the dynamics of states changing for more-easily-constructed energy landscape mapping and fixed-point structure construction. When the energy landscape can accept changes over time, the bifurcation will become accessible, which will further illustrate the process of attractors becoming deeper and farther away and suggest intervention methods. Our conclusion on behavior aims to indicate that neurons may not affect the behavior itself, but rather their range or possible dynamic (Fig. 4-5), which can be seen as not behavior continuous latent states, but the potential energy field are affected. In the neuron part, as mentioned by previous researchers[27], we also concern about whether the noise or input projected from other nuclei drive the neuron directly, or the neural structure drives itself autonomously but the noise or input changes the potential energy field.

## Method

### Animal subjects

A total of 7 adult male mice (C57BL/6J, 8 weeks old) were used in this study (Vital River Laboratory Animal Technology Co., Ltd.). All experiments were approved by Experimental Animal Welfare Ethics Review Committee, Zhejiang University (Ethical batch number, ZJU20220290). Mice were housed in individually ventilated cages under a normal 12 h light/12 h dark cycle, temperature was maintained between 19 °C and 23 °C and humidity between 50% and 65%.

### Viral injection

We used adeno-associated virus (AAV) expressing GCaMP6s, a genetic encoded fluorescence calcium indicator [1], to label M2 cortex principal neurons. Mice were anaesthetized with 1∼2% isoflurane in oxygen at a flow rate of 0.4 liter/min and mounted on a stereotactic frame (Model 68801, RWD Life Sciences, Inc., Shenzhen, China), while mice body temperature was heated on a heating pad. Sterile ocular lubricant ointment was applied to mouse corneas to prevent drying. Mouse scalp fur was shaved, and the skin was cleaned with tincture of iodine and 75% alcohol three times. A non-invasive surgery technique was used to minimize tissue damage and dura overgrowth underneath the optical window[40]. In a brief, a 5-mm-diameter hole was drilled through the right side, and the skull carefully removed (center: 1.5 mm anterior, 0.5 mm lateral to Bregma), using a high-speed rotary micro drill (Model 78001, RWD Life Sciences, Inc., Shenzhen, China). After skull debris were removed, a pair of fine forceps was used to remove 1 mm^2^ of dura around the injection area. The glass pipette tip touched the intact pia mater and injected AAV2/9-mCaMKIIa-GCaMP6s (500 nl, ∼2E+12V.G./mL, Shanghai Taitool Bioscience Co., Ltd). Then, a glass slide substitute was fixed with bio glue at the location where the skull was removed. The scalp was disinfected with iodine again and sutured until it was reopened two weeks later.

### Motor learning task

All mice were acclimated to the head-mounted miniScope prior to the experiment. The mice underwent a five-day training regimen, with each session preceded by a 5-minute test. Before the test, all mice familiarize with the environment for 1 hour. At the beginning of the test, the rotating rod was set to a slow speed of 4 rpm, and the mice were gently placed on the rod while their behavior was recorded using a camera. Filming was ceased either when the mice fell off the rod or when the 5-minute test duration had elapsed. Following the test, the mice proceeded to the training phase, during which the rotation speed of the rod was gradually increased from 4 rpm to 40 rpm over the course of 100 seconds[41]. If a mouse fell off during training, it was assisted back onto the rod to continue the session. Each day’s training was repeated 20 times to help the mice learn to stay on the rotating rod. The rotarod performance is defined as the maximum tolerable rpm each trail.

### Miniaturized microscope

A custom-built one-photon miniaturized fluorescence microscope (1P-miniFM) [42] was used to image fluorescence at selectable frequencies from 20Hz. 1P-miniFM weighs 2.9 g and has 1250 μm x 1050 μm maximum field of view on a cellular spatial resolution, with a wide range of 400 μm z-axis scanning. After achieving the in-focus position for the entire field of view, the base was fixed on the skull using dental cement. 1P-miniFM body is attached to mouse head on the base during the testing state and be detached during the training state.

### Behavioral video analysis

#### Pose estimation (DeepLabCut)

We utilized the behavioral tracking software DeepLabCut[21] to monitor the movements of mice during motor learning tasks. Eight anatomical landmarks were selected for tracking: the back, tail, left and right legs, left and right feet, and left and right toes. These landmarks were manually annotated in a subset of the behavioral video frames, which were then used to train a DeepLabCut model. The trained model was subsequently applied to all video recordings from the experiment, providing estimated positions for each tracked body part across every frame. To further enhance the quality of the landmark annotations and the DeepLabCut model, we conducted an accuracy check of the markers, extracted outlier frames for re-annotation, and improved the model accordingly.

#### Behavioral feature extraction

Mouse pose data, represented as X and Y pixel coordinates, was transformed into both internal and external features. Internal features were derived by calculating the point-to-point distances between body parts within each frame (N = 28 with 8 body parts) and combining these with the distances between each body part and the corresponding positions of all body parts in the subsequent frame (N = 64 with 8 body parts)[43]. This approach allowed us to capture detailed internal feature information.

To extract external features, an external anchor point was first determined. Specifically, we identified the maximum and minimum x and y coordinates for each body part across the entire time period. These coordinates represent the outermost boundaries of the bounding box that encompasses all body parts. The external anchor point was then defined by calculating the mean of these maximum and minimum values, which essentially provides the midpoint along each dimension (x and y), forming the geometric center of the bounding box. The coordinates of this center were concatenated to obtain the central point of the area. The distance from each body part to this external anchor point was calculated to represent the external positional features (N = 8 with 8 body parts). Additionally, the difference in the distance from each body part to the external anchor point between consecutive frames was used to represent the external velocity features (N = 8 with 8 body parts).

Given the presence of significant high-frequency noise when using DeepLabCut (DLC) for body part estimation, the extracted feature data were subjected to correction. Initially, a 10-frame median filter is applied across the data to mitigate the influence of outliers by replacing each value with the median of its neighboring points within a specified window. Following this, a low-pass filter with a 5 Hz cutoff frequency is employed to smooth the data by attenuating high-frequency noise, thus preserving the relevant low-frequency signals. Lastly, the processed data undergoes z-score normalization, which standardizes the data by scaling it to have a mean of zero and a standard deviation of one, facilitating comparability across different features. From this, we obtained 108 dimensions of behavioral characteristic data.

#### Dimensionality reduction

Non-linear dimensionality reduction techniques are instrumental in identifying sets of activity patterns that reside within a low-dimensional manifold embedded in the high-dimensional space of the data, even when this manifold is non-linear in nature. To achieve this, we employed Laplacian Eigenmaps (LEM) as our chosen method for non-linear dimensionality reduction. These techniques typically leverage the local relationships between adjacent data points to reconstruct a global distance metric. Given K data points in an N-dimensional space (where K is the number of frames and N is the number of behavioral features), a weighted graph was constructed with K nodes and edges linking neighboring data points. The edge weights were based on the Euclidean distances between points. A Laplacian eigenmap was then applied for dimensionality reduction, preserving local relationships by embedding the data into a lower-dimensional space using the eigenvectors associated with the smallest non-zero eigenvalues. This approach effectively captured the data’s intrinsic geometry while reducing complexity. For subsequent analysis, we focused on the six leading eigenvectors.

#### Analysis of behavior stabilization

We assessed the stabilization of operant behavior sequences by analyzing the trajectory of each mouse across trials[22]. For each mouse, 30-time intervals corresponding to the peaks and valleys of the trajectory from PC1 were recorded. Pairwise distances between these trajectories from PC1 within the defined time intervals were computed using dynamic time warping (DTW) with the MATLAB function “DTW”. The median trial-to-trial trajectory distance was then calculated for each mouse. These median distances were averaged across all mice for each training day.

#### Calcium imaging data processing

Images were firstly manually cut to a consistent viewing field. Further processing was performed using the MiniAn pipeline[44], including Denoising, Background removal, Motion correction, Initialization, and Spatial and temporal updates. In a brief, it is a pipeline enhancing background removal for miniScope images, including rigid motion correction using the NormCorre algorithm and source extraction using the CNMF-E algorithm. We rejected spurious ‘neurons’ resulting from imperfect motion correction by calculating the frame-by-frame correlation between a candidate’s activity trace and the magnitude of the motion correction vector and rejecting candidates with a correlation of 0.8 or greater while distance less than 10 pixel. Calcium dynamics are passed through a low-pass filter, and then simulated by convolving the spikes with a temporal kernel:

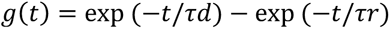

with rise time *τr* = 5 frames and decay time *τd* = 60 frames (20hz). Total of 334 ± 98 neurons from 4 mice in 5 days were detected.

#### Cross-registration

To register cells across sessions, the MiniAn cross-registration pipeline was used to align the field of view[44]. A standard cross-correlation based on a template-matching algorithm was used to estimate the translational shifts for each session relative to the template and then correct for this shift. In a brief, a pair of closest cells to each other, less than 5 pixels apart, is considered as the same cell across sessions.

#### Dynamical system models of neural data

The neural data consist of all z-scored cell signals. We model neural activity using a recurrent switching linear dynamical system (rSLDS)[27, 45, 46]. In a brief, rSLDS is a generative model, which consists of three systems: emission system, dynamical system, and transitional system.

The neural data is considered as observation *y*, while a set of continuous latent factors (*x*) that captures the low-dimensional nature of neural activity.

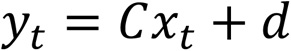

The activity of recorded neurons is recovered by modelling activity as a linear noisy Gaussian observation *y*_*t*_ ∈ ℝ^*N*^ where *N* is the number of recorded neurons. For *C* ∈ ℝ^*N*×*D*^ and *d*∼*N*(0, *S*), a gaussian random variable. These consist of emission system.

At any time t, continuous latent factors (*x*) are in a linear dynamical system, which means it has a high probability of moving in a consistent direction (over a period) of enlargement, contraction, saddle orientation, or (and) rotation. This consistent dynamic characteristic over a period is discrete state *z_t_*:

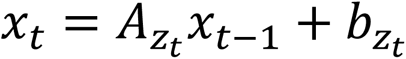

where *A*_*k*_ ∈ ℝ^*d*×*d*^ is a dynamics matrix, and *b*_*k*_ ∈ ℝ^*d*^ is a bias vector, where *d* is the dimensionality of the latent space. Specially, it is inferred that fixed points in discrete states will not move over time. Fixed point is the center of the dynamics system:

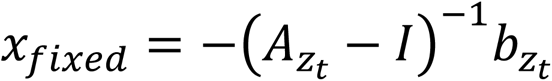

where *I* is the diagonal matrix in *d* × *d*. While there is only one global *z_t_*, it is a Simple Linear Dynamical System (LDS). While the discrete state *z_t_* transition follows a fixed Markov transition matrix, it is a Switching Linear Dynamical Systems (SLDS). However, rSLDS allows for the transitions between states to depend recurrently on the continuous latent factors (*x*) as follows:

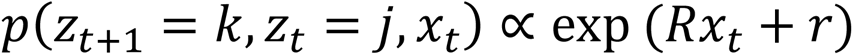

where *R* and *r* parameterizes a map from the previous discrete state and continuous state using a SoftMax link function to a distribution over the next discrete states, which means the space of continuous latent factors (*x*) is roughly divided into different regions with different dynamic structures *z_t_*.

Overall, the system parameters that rSLDS needs to learn:

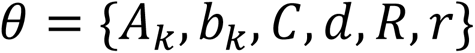

These parameters are estimated using maximum likelihood using approximate variational inference methods as described in detail in[45, 46].

Model performance is reported as the evidence lower bound (ELBO) which is equivalent to the Kullback-Leibler divergence between the approximate and true posterior. Considering the computational pressure caused by excessively high dimensions and the ELBO performance of different states of *z_t_* (Supplementary Fig. 2C) and dimension (*d*) (Supplementary Fig. 2D), three states of *z_t_* and six dimensions (*d*) are used.

#### Distance between attractors

The transformation matrix *T*_*k*_ reflects the vector of changes in continuous latent factors (*x*) when it is in a discrete state *z_t_*:

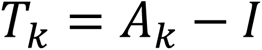

Where *A*_*k*_ ∈ ℝ^*d*×*d*^ is the dynamics matrix of *k* and *I* is the diagonal matrix in *d* × *d*. The enlargement, contraction, saddle orientation, or (and) rotation characteristics are determined by eigenvalues *λ*_*a*_ of the *T*_*k*_.

The sum of eigenvalues is equal to the trace of the transition matrix, which is the sum of the diagonal elements of the matrix. Trace has the property of maintaining certain dynamic characteristics. The values of the determinant of a matrix are the product of the eigenvalues. Traces can reflect the stability characteristics of the system. If the trace is negative while all eigenvalues are negative, the dynamic system is inferred stable (Supplementary Fig. 2F). The discrete state *z_t_* which has stable dynamic system with the fixed point within the region is considered as attractor.

#### Estimation of time constants

We use the eigenvalue *λ*_*a*_ of the dynamic matrix of the system to estimate the time constants *τ*_*a*_ of each mode of the linear dynamic system:

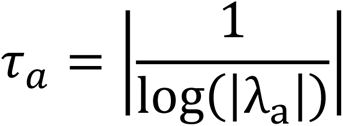

#### Low dimensional (PCA) representation of non-linear system

To reduce the dimensionality of our behavioral and neural data and capture the most significant variance, we employed Principal Component Analysis (PCA). PCA is particularly well-suited for this task as it enables us to describe our high-dimensional, non-linear system in a more concise and interpretable manner by projecting the data onto a lower-dimensional space.

The data was first standardized to ensure equal contribution from each feature. Next, the covariance matrix was computed to reveal the relationships between variables. Eigenvalues and eigenvectors were then derived from the covariance matrix, representing the magnitude and direction of the principal components, respectively. The principal components were ranked according to their eigenvalues, and the top components, capturing the majority of variance, were selected for further analysis.

Particularly, since the latent states are invariant to linear transformations, it is possible to apply a suitable transformation of PCA for both continuous latent factors (*x*) and its dynamics matrix (*A*). For our analysis, we retained the first two principal components (PC1 and PC2) as they accounted for the most significant portion of the variance (60.4% ± 2.4%), providing a reduced yet informative representation of the original high-dimensional system.

#### Autocorrelation

Recent studies indicate that the temporal stability of neuronal activity can be assessed by examining the structure of the neuronal autocorrelation in neuronal populations[29]. Meanwhile, the autocorrelation analysis is also used to detect the periodicity of signals[24]. We computed the autocorrelation structure for PC1 of neuronal space and behavioral space as a function of time lag and estimated its decoupling time or periodicity.

The result of autocorrelation can be interpreted as the correlation of a signal with a delayed copy of itself as a function of delay.

Given a real signal *f*(*t*), the continuous autocorrelation *R*_*ff*_(*τ*) is most often defined as the expected value of product of *f*(*t*) with itself, at lag *τ*:

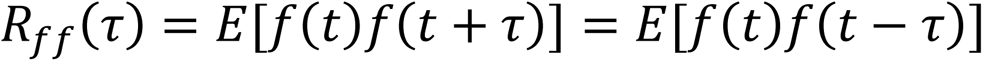

Where −∞ < *t* < ∞, and *E* is the expected value operator.

In general, *f*(*t*) is a discrete sequence, and computes raw correlations with no normalization:

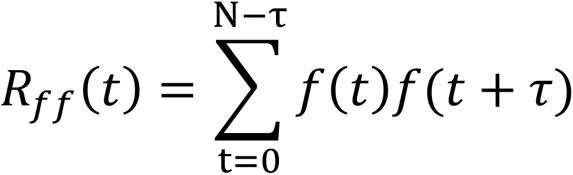

Decoupling time indicates the time lag from central peak to the time bin when the coefficient reaches zero for the first time, which means the shortest time *τ* for *f*(*t*) to be orthogonal to *f* (*t* + *τ*).

Periodicity is inferred from the relative height, measured by the height of the peak itself and its position relative to other peaks, of the first non-central peak of autocorrelation.

#### Energy landscapes

The arrow in neural PCA space is from latent factors at time *x*_*t*_ to inferred latent factors at time

*x*_*t*+1_, which means the force *f_t_* at time *t* can be inferred as follows:

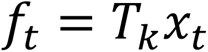

Where the transformation matrix *T*_*k*_ is mentioned above. To construct the energy landscape, we first defined a grid over the region of interest, which generated matrices X and Y representing the coordinates of grid points. For each point on the grid, we computed its distance to all other points using the Euclidean distance formula. Subsequently, the energy at each grid point was then calculated as logarithm of the integral of the force magnitudes over the distance. This methodology allows for the accurate mapping of the energy landscape based on spatial and force data, facilitating a comprehensive understanding of the system’s energy distribution.

#### Flood Fill Algorithm

To calculate the relative depth of attractors, we introduce flood fill algorithm to create a valley-filled landscape to measure the depth by the residual of two landscape. Flood fill algorithm is mainly accomplished by “imfill” function implemented in MATLAB. We firstly use “imregionalmin” function in MATLAB to determine all the valley in landscape, and imfill function will fill all the holes in field. Based on this, we get two different energy fields, whose residual is the relative depth we require.

#### Identification of Behavioral Place Cells

Calcium imaging data, along with corresponding position data, was employed to identify “place cell” neurons that display location-specific firing patterns. The high-dimensional position data, initially comprising six spatial dimensions (data after nonlinear dimensionality reduction by LEM), was reduced to two principal components through Principal Component Analysis (PCA) to facilitate the spatial mapping of neural activity. Concurrently, the calcium imaging data was standardized across neurons by applying z-scoring, which normalized the neuronal activity by centering the data and scaling it to unit variance.

Subsequently, the animal’s spatial environment was divided into a 10×10 grid, allowing for the computation of average calcium activity within each spatial bin. This process yielded a firing rate map for each neuron, reflecting the neuron’s activity across different spatial locations. The correlation between the firing rate map of each neuron and the animal’s spatial position was then computed. Neurons exhibiting a correlation coefficient greater than 0.50 were identified as place cells, indicating a significant relationship between the neuron’s activity and the animal’s location in the environment.

#### Behavioral variability

Behavioral variability is evaluated by using the sliding variance of behavior. Given a potential periodicity of approximately 2 seconds in main behavioral components, a 2.5-seconds sliding window is used to calculate the sliding variance of each point, which means sliding variance indicates the variance of PC1 within 2.5 seconds after this moment.

#### Predicting choice from neural population

We used logistic regression[47, 48] to estimate how the weighted sum of neural activity predicted the main behavioral components (behavioral space PC1) or behavioral variability. The model fits each recording session separately as described above using the glm-net package in MATLAB, using a nested fivefold cross-validation. To quantify the predictive power, we computed the Pearson’s correlation coefficient (PCC) between behavioral components or variability predicted by the model in the test set and the corresponding real components or variability.

#### Quantification and statistical analysis

Data were processed and analyzed using Python, MATLAB, and GraphPad (GraphPad PRISM 11). All data were analyzed using two-tailed non-parametric tests. Friedman test was used for paired samples. Kolmogorov-Smirnov test was used for non-paired samples. Multiple comparisons were corrected with Dunn’s multiple comparisons correction. Not significant (NS), P > 0.05; *P < 0.05; **P < 0.01; ***P < 0.001.

---

**Supplementary Figure 1.**
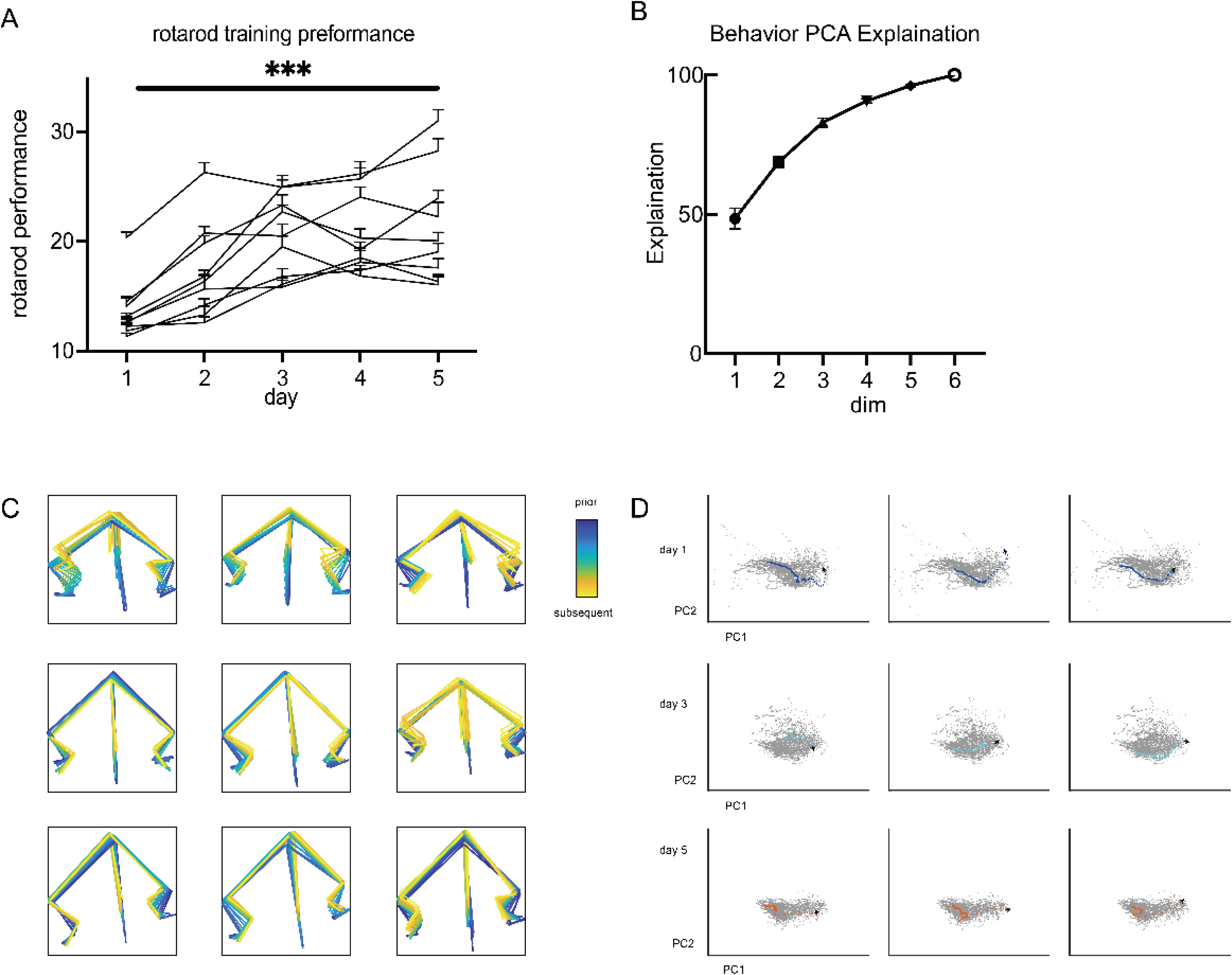
(A) Rotarod performance across the training days (p < 0.001, n = 7 mice, paired t-test), showing a gradual improvement in the behavior of the mice. Mouse’s performance was scored by time latency of rotarod task. (B) Behavior variance explained by PCA analysis. (C) Superposition of the mice’s behavioral posture estimation over each step of day 1,3,5 in real world, with a consistent trajectory shown in D. (D) Trajectories in low-dimensional space of mice’s steps of day 1,3,5 behavior space for a single mouse. Three steps’ trajectories each day are shown corresponding to C.

**Supplementary Figure 2.**
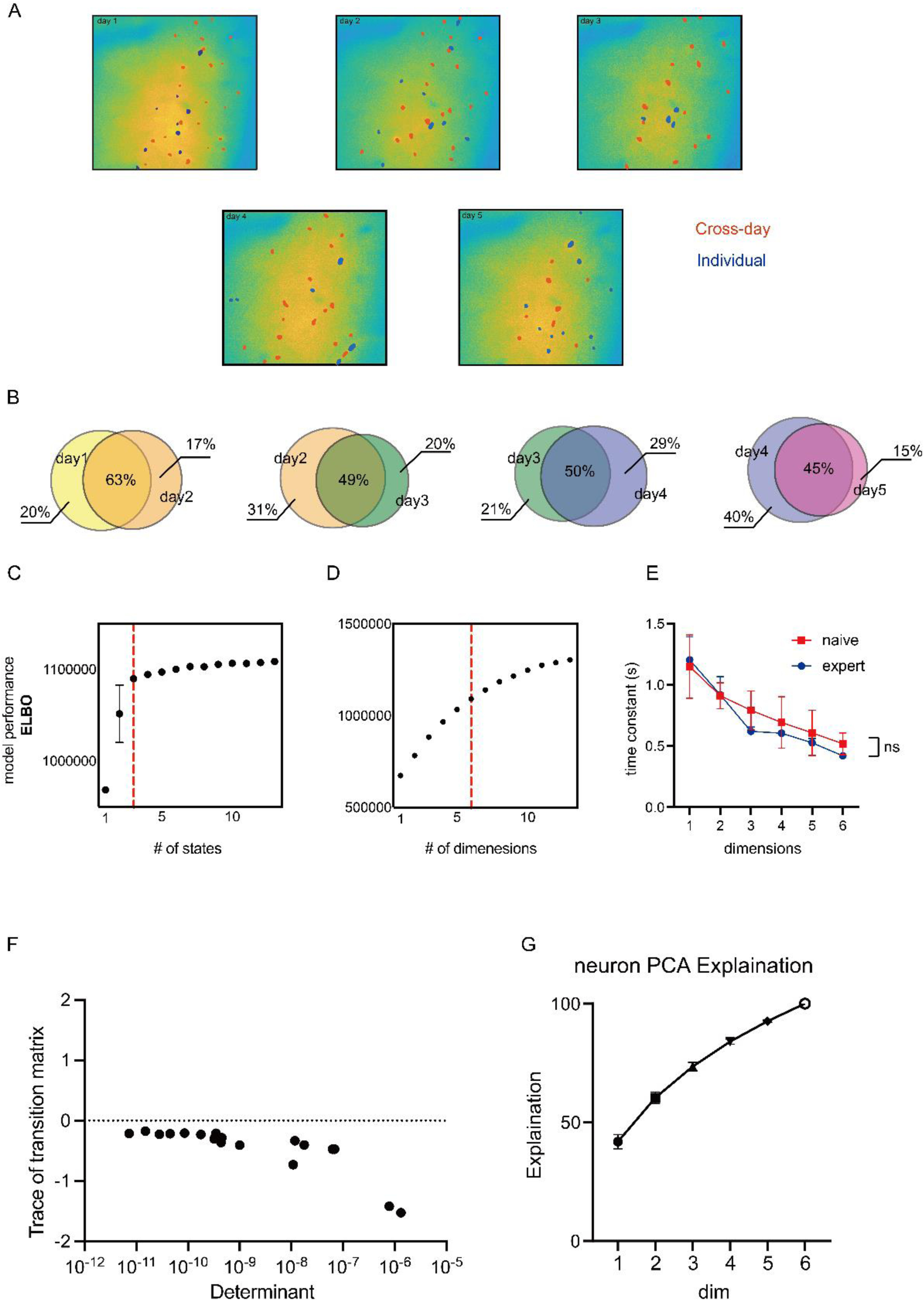
(A) Distribution of firing cells in the field of view. Cells firing across the day were marked in red, others firing on single day were marked in blue. (B) Venn diagram of the proportion of cells fired across the day in all cells observed in the field of view on the two preceding days. (C) Left, relationship between model performance and states, showing a sharp upward trend. Right, relationship between model performance and dimensions. Model performance is measured as the evidence lower bound (ELBO), which is equivalent to the Kullback-Leibler divergence between the approximate and true posterior. Red line represents the chosen parament. (D) Curves of the time constants of linear dynamical systems for expert mice (triangles) versus naive mice (squares) in different dimensions. (E) The determinant and trace for each primary attractor matrix. All transition matrixes stay in the fourth quadrant. The determinant is represented logarithmically on the X-axis. (F) Neural variance explained by PCA analysis.

