## Supplementary figures and images for "Separated attractors in neural landscape of motor cortex encoding motor learning"

### supplementary figure_1.png

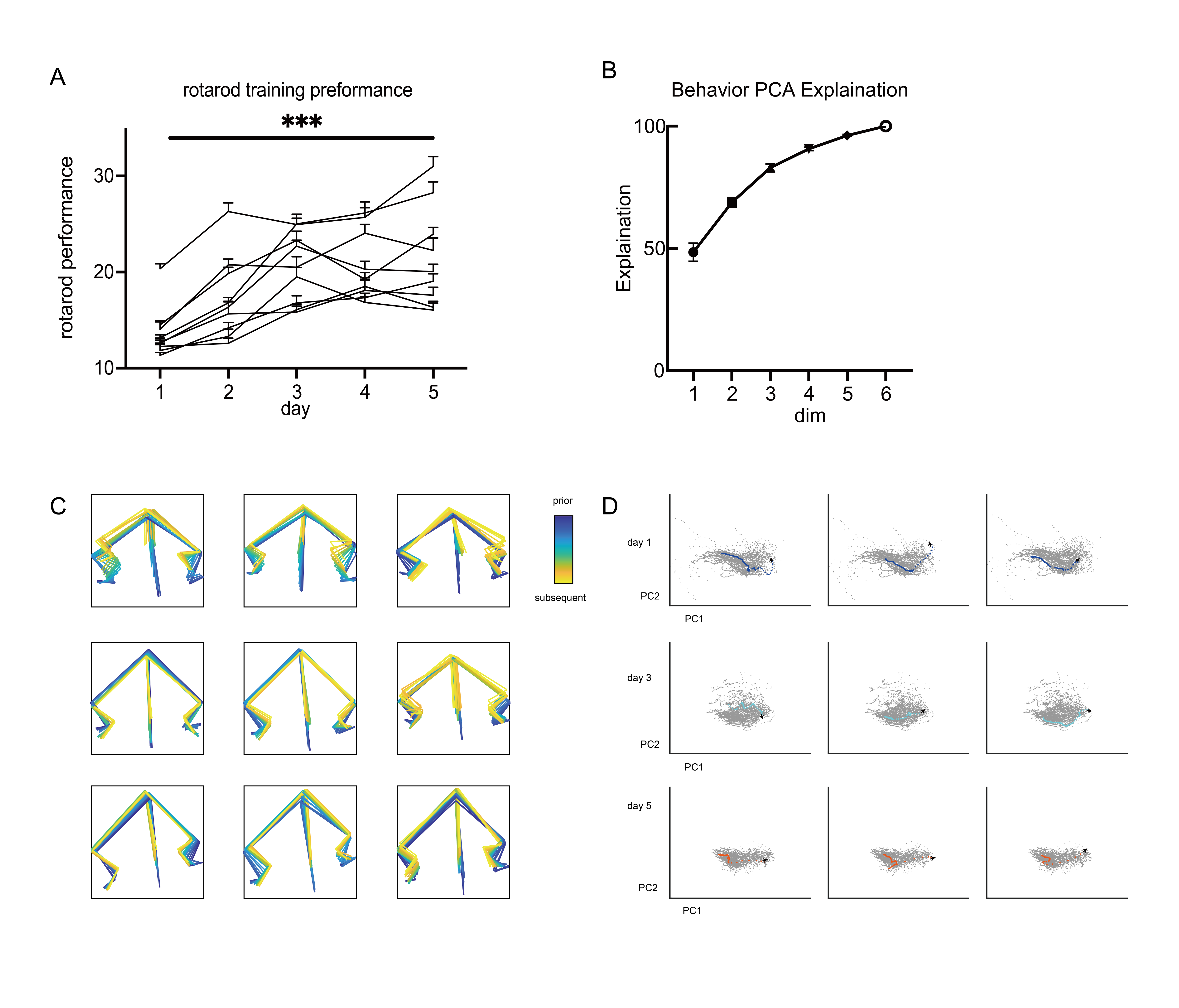

### supplementary figure_2.png

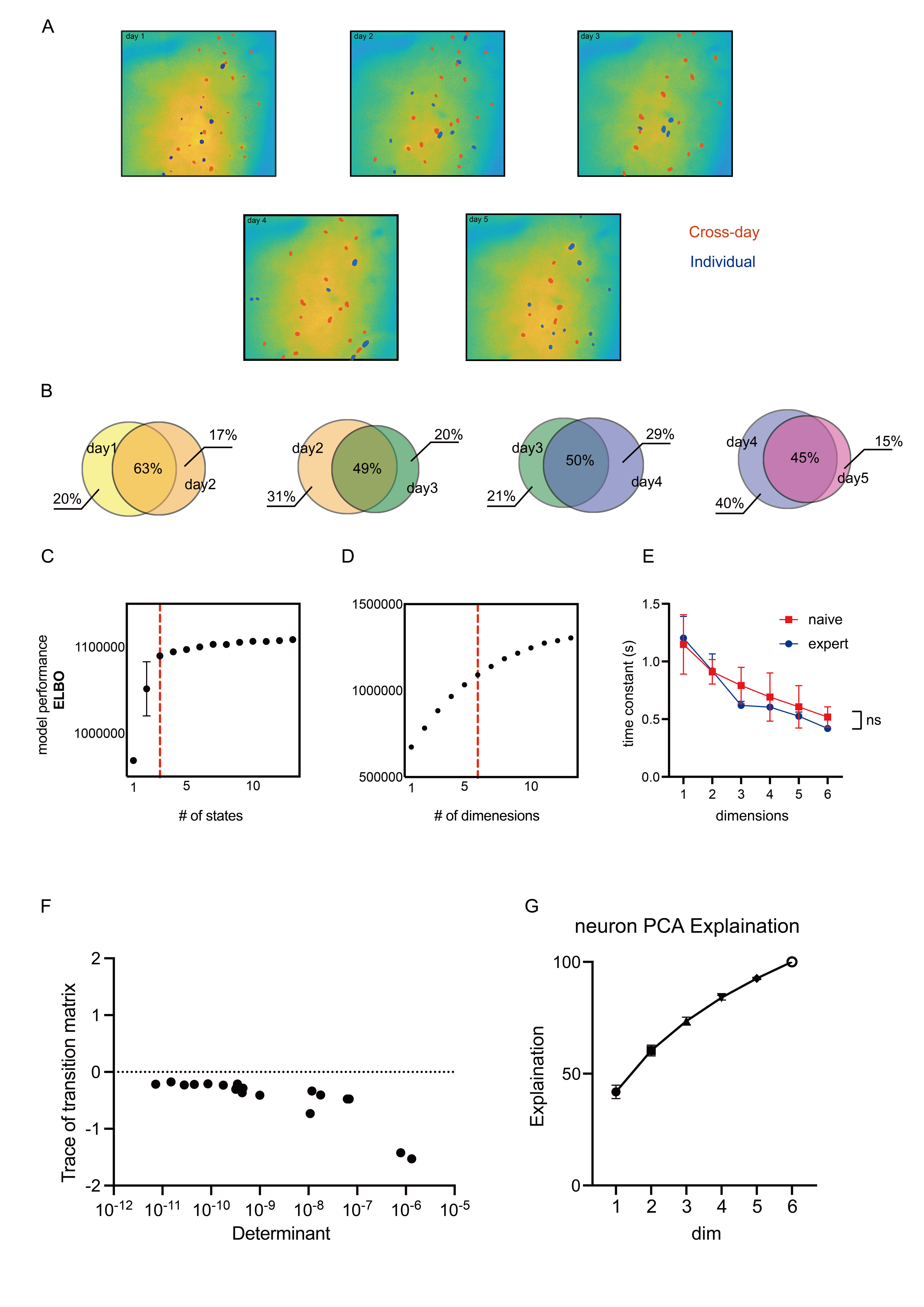
